# Alternative splicing expands the antiviral IFITM repertoire in Chinese horseshoe bats

**DOI:** 10.1101/2023.12.04.569605

**Authors:** Nelly Mak, Dan Zhang, Xiaomeng Li, Kazi Rahman, Siddhartha A.K. Datta, Jordan Taylor, Jingyan Liu, Zhengli Shi, Nigel Temperton, Aaron T. Irving, Alex A. Compton, Richard D. Sloan

**Affiliations:** Centre for Inflammation Research, Institute for Regeneration and Repair, University of Edinburgh, Edinburgh, UK; Deanery of Biomedical Sciences, Edinburgh Medical School, University of Edinburgh, Edinburgh, UK; Zhejiang University-University of Edinburgh Institute, Zhejiang University School of Medicine, Zhejiang University, Haining, China; HIV Dynamics and Replication Program, National Cancer Institute, National Institutes of Health, Frederick, MD, USA; CAS Key Laboratory of Special Pathogens and Biosafety, Wuhan Institute of Virology, Chinese Academy of Sciences, Wuhan, China; Viral Pseudotype Unit, Medway School of Pharmacy, University of Kent and Greenwich Chatham Maritime, Kent, UK; Second Affiliated Hospital, School of Medicine, Zhejiang University, Hangzhou, China; Centre for Infection, Immunity & Cancer, Zhejiang University-University of Edinburgh Institute, Haining, China; College of Medicine and Veterinary Medicine, University of Edinburgh, Edinburgh, United Kingdom; Biomedical and Health Translational Research Centre of Zhejiang Province, Haining, China

## Abstract

The interferon response is shaped by the evolutionary arms race between hosts and the pathogens they carry. The human interferon-induced transmembrane protein (IFITM) family consists of three antiviral *IFITM* genes that arose by gene duplication, they restrict virus entry and are key players of the interferon response. Yet, little is known about IFITMs in other mammals. Here, we identified an *IFITM* gene in Chinese horseshoe bat, a natural host of SARS-coronaviruses, that is alternatively spliced to produce two IFITM isoforms. These bat IFITMs have conserved structures in vitro and differential antiviral activities against influenza A virus and coronaviruses including SARS- and MERS-coronavirus. In parallel with human IFITM1-3, the bat IFITM isoforms localize to distinct cellular compartments. Further analysis of IFITM repertoires in 205 mammals reveals that alternative splicing is a ubiquitous strategy for IFITM diversification, albeit less widely adopted than gene duplication. These findings showcase an example of convergent evolution where species-specific selection pressures led to expansion of the IFITM family through multiple means, underscoring the importance of IFITM diversity as a component of innate immunity.

## Introduction

Human and other animals possess interferon stimulated genes (ISGs) that encode antiviral proteins acting as the first line of defense against virus infections. Interferon-induced transmembrane proteins (IFITMs) are a family of antiviral proteins that inhibit virus entry into target cells and are upregulated by type I, II and III interferons (IFN). The human IFITM family consists of three antiviral IFITMs with high sequence similarity (IFITM1, IFITM2 and IFITM3) and two IFITMs that are not interferon-inducible and not known to be involved in immunity (IFITM5 and IFITM10) (Zhang *et al*, 2012; Zhao *et al*, 2019). IFITM3 is the most well-studied owing to its potency against Influenza A virus (IAV) and many other viruses, such as HIV-1 and Dengue virus (Brass *et al*, 2009; Lu *et al*, 2011). The effect of IFITMs on coronaviruses (CoVs) is less clear cut. While CoVs are generally inhibited by overexpression of IFITMs, endogenous IFITMs have little effect, or may even promote SARS-CoV-1 and SARS-CoV-2 infection (Shi *et al*, 2020; Xie *et al*, 2023a). An exception is human coronavirus OC43, which is always enhanced by IFITM2 and IFITM3 (Zhao *et al*, 2014; Zhao *et al*, 2017). Beyond their antiviral activity, IFITMs have pleiotropic effects such as regulating interferon production, adaptive T- and B-cell responses, and may even influence cancer growth (Gomez-Herranz *et al*, 2023; Rajapaksa *et al*, 2020).

IFITMs inhibit virus-cell membrane fusion by mechanisms that are not fully understood. The best current working model suggests that the IFITM3 amphipathic helix inserts into membranes to induce a negative membrane curvature that disfavors the formation of a fusion pore, a critical step in membrane fusion (Chesarino *et al*, 2017; Desai *et al*, 2014; Guo *et al*, 2021; Li *et al*, 2013). Cholesterol binding was recently shown to be crucial for IFITM3 antiviral activity (Das *et al*, 2022; Rahman *et al*, 2022). However, the role played by cholesterol in IFITM-mediated inhibition of virus entry is less straightforward with several alternative mechanisms being proposed (Amini-Bavil-Olyaee *et al*, 2013; Klein *et al*, 2023; Lin *et al*, 2013; Wrensch *et al*, 2014). IFITMs contain a canonical CD225 domain that is comprised of an intramembrane domain and a conserved intracellular loop, and it contains residues that can be post-translationally modified to alter IFITM function (John *et al*, 2013; Yount *et al*, 2012; Yount *et al*, 2010). The _20_YXXΦ_23_ motif in the N-terminal domain of IFITM2 and IFITM3 serves as an endocytic signal for their localization to endolysosome membranes, whereas the absence of this motif in IFITM1 results in its surface localization (Jia *et al*, 2014).

Since the discovery of IFITMs as antiviral proteins in 2009, efforts from numerous research groups have led to significant advances in our understanding of IFITM function. Yet, most studies have focused on human and mouse IFITMs, leaving a gap in our knowledge on IFITMs in other species. The wider ISG biology field faces the same problem, with increasing interest in examining immunity in non-model species given their epidemiological significance. Reservoir hosts such as birds, pigs and bats harbor zoonotic viruses with the potential to ‘jump’ into human, and these spillover events can spark virus outbreaks. In particular, bats have attracted immense research interest in light of recent virus outbreaks. Bats make up 21% of all mammalian species, they are most abundant in land with substantial human activity and likely harbor more zoonotic viruses compared to other mammals (Burgin *et al*, 2018; Gibb *et al*, 2020; Olival *et al*, 2017). Examples of bat-originated viruses that have caused epidemics or pandemics in the human population are Marburg virus, Nipah virus, and coronaviruses including MERS-CoV, SARS-CoVs and the common cold human coronavirus 229E (HCoV-229E) (Cui *et al*, 2019; Letko *et al*, 2020; Streicker & Gilbert, 2020). SARS-CoVs likely originated from horseshoe bats, with viruses closely related to SARS-CoV-1 and SARS-CoV-2 being detected in bats in the *Rhinolophidae* family (Lau *et al*, 2005; Temmam *et al*, 2022; Zhou *et al*, 2020). Many behavioral, ecological and molecular factors may predispose bats to acting as viral reservoirs, among which arguably the most important is their unique balance between host defense mechanisms and immune tolerance that may support their high metabolism (Gonzalez & Banerjee, 2022; Irving *et al*, 2021). Unfortunately, research on bat immunity is limited by the lack of reagents and the low availability of annotated bat genomes, with only two *Rhinolophus* genomes published in the NCBI RefSeq database at the time of this study (*R. ferrumequinum*, GCF_004115265.2; *R. sinicus*, GCF_001888835.1).

Host and viruses need to constantly evolve to gain a survival advantage over each other, viruses therefore impose strong selection pressures on the host immune system (Brockhurst *et al*, 2014). In fact, several non-human ISGs have been shown to have differential functional capacity compared to their human counterparts (Zhang & Irving, 2023). ISGs also have a higher rate of gene duplication compared to other genes which may enhance their potency and diversity (Shaw *et al*, 2017). Duplication of *IFITM* genes in a species-specific manner generates distinct IFITM repertoires (Compton *et al*, 2016; Zhang *et al*., 2012). Although several phylogenetic studies have examined *IFITM* genes in non-human species, a holistic analysis involving a larger subset of species is lacking. Previous studies also focused on the number and origin of *IFITM* genes in each species, without examining the transcripts they encode (Benfield *et al*, 2020; Hickford *et al*, 2012; Schelle *et al*, 2023). The latter is of interest because generation of alternatively spliced transcripts is a useful strategy in evolution to enhance transcriptome and proteome diversity (Keren *et al*, 2010). For example, alternative splicing generates an N-terminus truncated IFITM2 in human immune cells, which displays differential antiviral specificity against different HIV-1 strains (Wu *et al*, 2017). Alternative splicing is however a double-edged sword as it may be a source of pathology. The human IFITM3 rs12252-C allele is predicted to produce an alternatively spliced transcript that encodes an N-terminus truncated IFITM3, and has been associated with severe influenza, HIV-1 and COVID-19 (Everitt *et al*, 2012; Zhang *et al*, 2015; Zhang *et al*, 2020; Zhang *et al*, 2013). However, attempts to identify this mutant protein have thus far been unsuccessful (Alghamdi *et al*, 2021; Randolph *et al*, 2017).

Characterizing antiviral immunity in reservoir species helps address major questions: How evolutionarily conserved or distinct are ISGs under lineage-specific selection pressures? Do ISG repertoires shape species-specific zoonotic barriers? In this study, we examined the conservation of IFITM antiviral activity in bats using *R. sinicus* as a model owing to its role as the natural host of SARS-CoVs and the availability of sequencing data. We then further characterized IFITM repertoires in mammals more broadly to gain insights into how alternative splicing contribute to IFITM diversity.

## Results

### *R. sinicus* possesses an *IFITM* gene that encodes two IFITM splice variants

To characterize the IFITM repertoire in *R. sinicus*, we searched the available *R. sinicus* genome for homologs of human immune-related IFITM1-3. We identified a single gene, *LOC109436297*, that shows highest homology with *IFITM3* and is flanked by *B4GALNT4*, similar to the human *IFITM* locus **(Fig. 1A)**. Alternative first exon splicing of the gene generates two predicted protein-coding mRNA transcripts that are distinct at the N-terminus (XM_019714804.1 and XM_019714805.1). *R. sinicus* thus potentially expresses two IFITM isoforms which we referred to as rsIFITMa (XP_019570363.1) and rsIFITMb (XP_019570364.1), and collectively as rsIFITMs in this article. Orthologs of *IFITM5* and *IFITM10* are also present in the *R. sinicus* genome. They do not contain interferon-stimulated response elements (ISRE) around the transcription start site and like their human counterparts, are unlikely to be antiviral, thus were not examined in this study **(Fig. EV1A)**.

**Figure 1:**
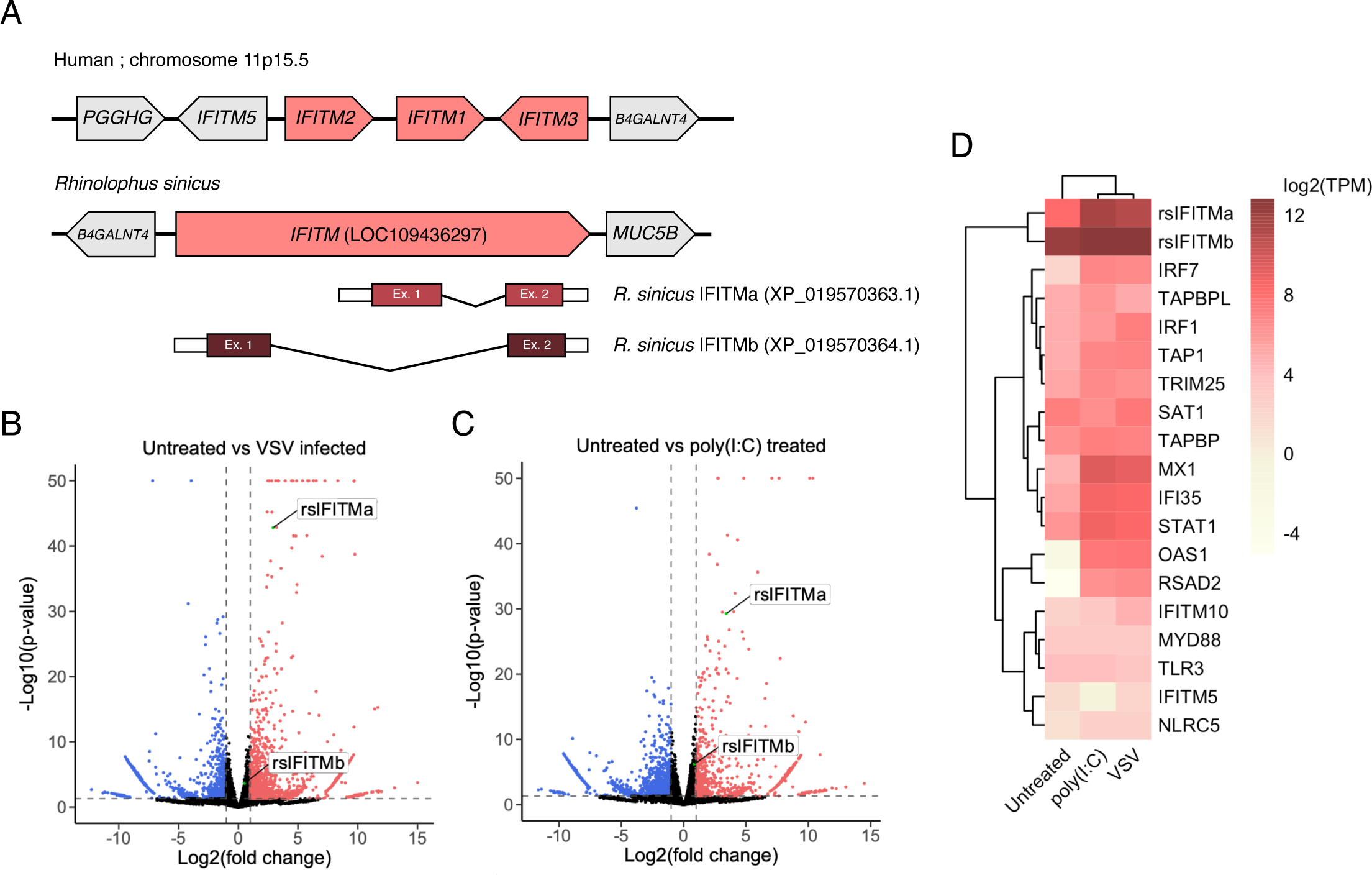
*Rhinolophus sinicus* possess an *IFITM* gene that encodes two distinct IFITM isoforms **A.** Schematic representation of the human *IFITM1-3* loci and *R. sinicus IFITM* (LOC109436297) with flanking genes. The *R. sinicus IFITM* gene was identified by BLAST and produces two transcripts, IFITMa (XP_019570363.1) and IFITMb (XP_019570364.1). **B-C.** Volcano plots of differential expression of gene transcripts comparing untreated cells versus poly(I:C)-treated (B) or VSV-infected (D) cells. Upregulated (red) and downregulated (blue) gene transcripts are indicated. Data points with - log10(p-value) above 50 are plotted at the y-axis upper limit. **D.** Heatmap showing normalized expression log2(TPM) of rsIFITM transcripts and selected ISGs for each condition. Gene-level TPMs were calculated as the sum of transcript-level TPMs for genes excluding *rsIFITM*. TPM; transcripts per million.

Pairwise amino acid sequence alignment of rsIFITMa and rsIFITMb indicates a 79.4% sequence identity, with most variation at the N-terminus **(Fig. EV1B)**. rsIFITMa contains the _20_YEML_23_ endocytic motif, while rsIFITMb resembles IFITM1 in that it has a truncated N-terminus lacking this motif. Alignment of human and *R. sinicus* IFITMs shows that 45 out of 50 residues in the canonical CD225 domain are conserved. Importantly, residues that undergo post-translational modifications such as ubiquitination (at Lys83, Lys88 and Lys104) and palmitoylation (at Cys71, Cys72 and Cys105) are conserved across these proteins, in line with a previous study that showed conservation of these residues in bat IFITMs (Benfield *et al*., 2020; Yount *et al*., 2012). However, Lys24, which was reported to be the most robustly ubiquitinated in IFITM3 is lost in rsIFITMb. Residues involved in IFITM oligomerization (Gly91 and Gly95) are also conserved (Rahman *et al*, 2020; Zhao *et al*., 2017).

To examine whether rsIFITMa and rsIFITMb are natively expressed, we performed RNA sequencing on *R. sinicus* kidney epithelial (RsKT.01) cells. Both rsIFITM isoforms were endogenously expressed and upregulated upon poly(I:C) treatment or infection with vesicular stomatitis virus (VSV), which elicits an interferon response. **(Fig. 1B-D)**. rsIFITMa was significantly upregulated upon treatment, while the fold-induction of rsIFITMb was minimal due to its high basal expression. Absolute quantification of expression was not possible due to biased handling of reads which mapped to sequences shared by both isoforms during analysis. These findings suggest that *R. sinicus* possesses an *rsIFITM* gene that encodes two IFITM splice variants with distinct N-terminal domains.

### *R. sinicus* IFITMs have a structurally and functionally conserved amphipathic helix

The human IFITM3 amphipathic helix is required for antiviral activity by binding cholesterol and increasing membrane order (Chesarino *et al*., 2017; Rahman *et al*., 2022). rsIFITMa and rsIFITMb share an identical amphipathic helix, which only differs from the human IFITM amphipathic helices at the last residue **(Fig. 2A)**. *In silico* analyses show that this amino acid substitution preserved the helical structure of rsIFITM amphipathic helix, while slightly reducing its mean hydrophobic moment and increasing its hydrophobicity, implying a reduced amphipathicity compared to that of human IFITM2 or IFITM3 (IFITM2/3) **(Fig EV2A)**. Wider analysis of mammalian IFITMs reveals that bat IFITMs have amphipathic helices that are less amphipathic compared to human and other mammals **(Fig EV2B)**. Circular dichroism spectroscopy of synthetic peptides confirms similar and substantial helical content (56.0 – 63.1%) between human and *R. sinicus* IFITM amphipathic helices **(Fig. 2B-C)**. Next, we tested whether the rsIFITM amphipathic helix retained the ability to bind cholesterol using a previously established fluorescence-based *in vitro* binding assay (Rahman *et al*., 2022). Peptide binding to the cholesterol analog NBD-cholesterol was measured by fluorescence intensity and polarization. All human and *R. sinicus* IFITM amphipathic helices exhibited cholesterol binding activity, with rsIFITM amphipathic helix binding cholesterol to a similar extent as the human IFITM2/3 amphipathic helix **(Fig. 2D, EV2C)**.

**Figure 2:**
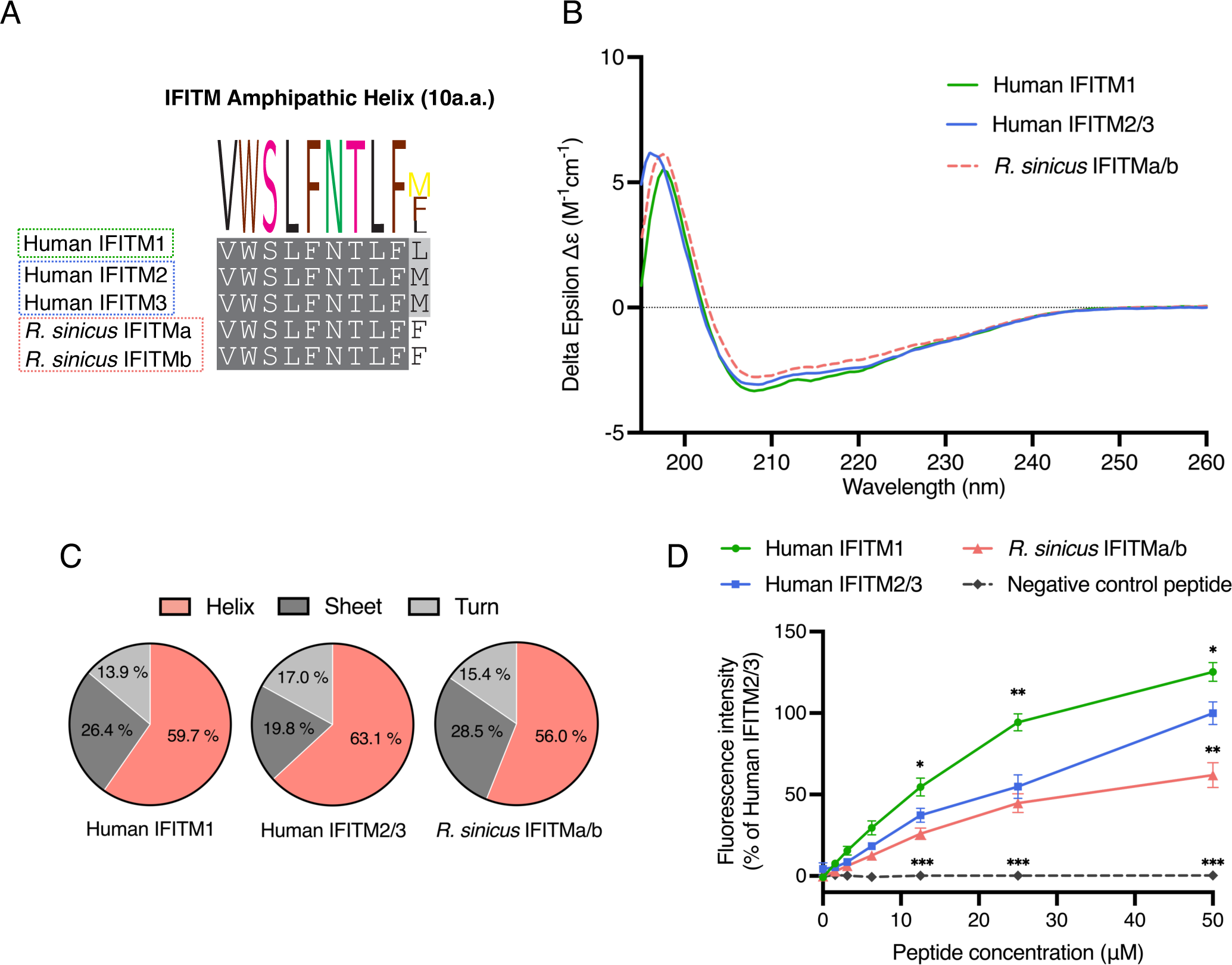
The amphipathic helix of *R. sinicus* IFITMs has conserved structure and function **A.** Protein sequence alignment of the amphipathic helix of immune-related human and *R. sinicus* IFITMs, with the consensus sequence shown as a sequence logo above. **B-C.** Structures of IFITM amphipathic helix peptides were characterized by circular dichroism spectroscopy to determine their secondary structure compositions. Spectra in (B) represent the average of six acquisitions. **D.** NBD-cholesterol fluorescence intensity was measured following incubation of IFITM amphipathic helix peptides (0-50μM) with NBD-cholesterol (500nM). Data points are normalised to 50μM human IFITM2/3. Error bars represent SEM of averages from 3 independent experiments. Statistical significance of difference between human IFITM2/3 and another peptide was determined by one-way ANOVA with Dunnett’s test; *p<0.05, **p<0.01, ***p<0.001.

To confirm that the rsIFITM amphipathic helix is sufficient to mediate IFITM antiviral activity, chimeric constructs were generated by substituting the amphipathic helix of human IFITM3 with that of rsIFITMs (IFITM3-AH [*R. sinicus*]) **(Fig EV2D)**. Expression of IFITM3 or IFITM3-AH [*R. sinicus*] led to potent inhibition of IAV infection, with no significant difference between the extent of inhibition **(Figure EV2E-F)**. Taken together, we show that the rsIFITM amphipathic helix has conserved secondary structure and function, albeit containing a mutation in the last amino acid.

### *R. sinicus* IFITM isoforms exhibit differential antiviral activity

To examine the antiviral activity of rsIFITMs, we expressed FLAG-tagged IFITM constructs in HEK293T cells and challenged them with IAV. Both rsIFITMs were well-expressed upon transfection, but at a lower level than human IFITMs **(Fig. 3A)**. To mitigate variation arising from inconsistent transfection efficiencies, transfected cells were stained and gated by flow cytometry to select only FLAG-positive cells for downstream analysis. Expression of human IFITM3 led to the strongest inhibition of IAV infection by 14-fold **(Fig. 3B, EV3A)**. Inhibition of IAV infection by human IFITM1 and IFITM2 were significant but less pronounced, which is consistent with previous studies (Brass *et al*., 2009). Both rsIFITMs inhibited IAV infection, the antiviral potency of rsIFITMa was comparable to IFITM3 and significantly greater than rsIFITMb (24-fold vs 3-fold inhibition, p=0.002).

**Figure 3:**
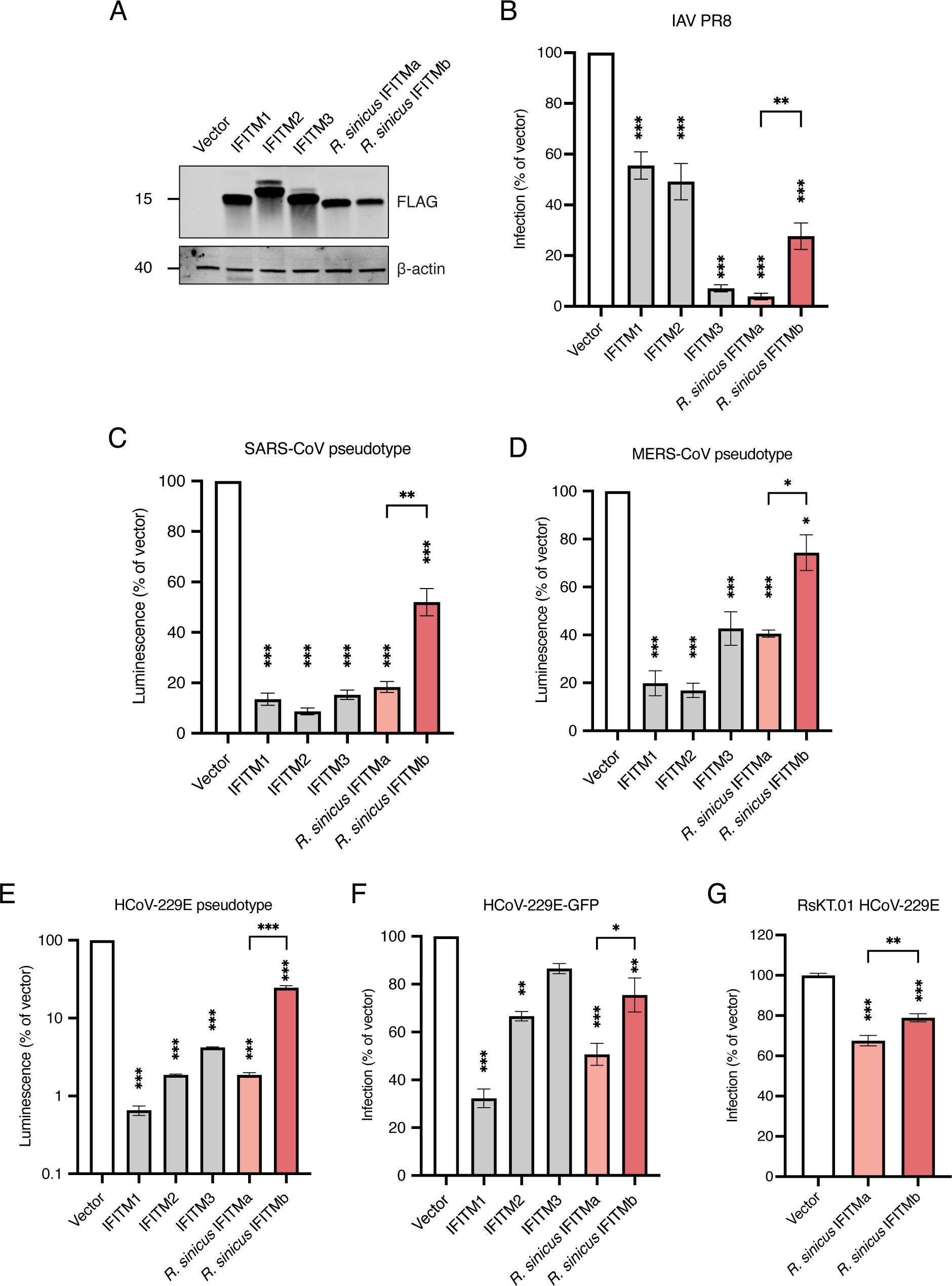
*R. sinicus* IFITMs exhibit differential antiviral activity against IAV and coronaviruses **A.** HEK293T cells were transfected with the indicated FLAG-tagged IFITMs. IFITM expression was detected by western blotting at 24 hours post-transfection. **B.** Transfected HEK293T cells were infected with IAV and analysed by flow cytometry at 18 hours post-infection. Error bars represent SEM of averages from 3 independent experiments, each performed in duplicate. **C-E.** HEK293T cells were co-transfected with the indicated IFITMs and coronavirus entry receptor (ACE2, DPP4 or APN) then transduced with SARS-CoV-1 (C), MERS-CoV (D) or HCoV-229E (E) pseudotypes respectively. Cells were lysed and analyzed by luciferase assay after 48 hours. Error bars represent SEM of averages from 3 independent experiments, each performed in at least three replicates. **F.** Huh7.5 cells were transfected with the indicated IFITMs and infected with GFP-tagged HCoV-229E virus (MOI=0.05). Cells were analysed by flow cytometry at 18 hours post-infection. Error bars represent SEM of averages from 3 independent experiments, each performed in duplicate. **G.** RsKT.01 cells stably expressing IFITMs were infected with serially diluted HCoV-229E virus and infection was quantified by TCID50 assay. Statistical significance of difference between vector- and IFITM-expressing cells were determined by one-way ANOVA with Dunnett’s test; Statistical significance of difference between *R. sinicus* IFITMa- and IFITMb-expressing cells were determined by unpaired t-test; *p<0.05, **p<0.01, ***p<0.001.

To further test whether the two rsIFITM isoforms also restrict coronaviruses to different extents, lentiviral-based pseudotyped viruses expressing spike protein from SARS-CoV-1, MERS-CoV or the HCoV-229E were used. Expression of human IFITM1-3 inhibited the entry of all three CoV pseudotypes in line with previous reports **(Fig. 3C-E)** (Huang *et al*, 2011; Wrensch *et al*., 2014). Both rsIFITM isoforms inhibited the CoV pseudotypes, and again, rsIFITMa showed a stronger restriction than rsIFITMb. IFITM-mediated inhibition of HCoV-229E infection was confirmed with replication-competent HCoV-229E-GFP, which shows a similar pattern of inhibition by rsIFITMa and rsIFITMb **(Fig. 3F, EV3B)**. Next, we sought to confirm that rsIFITMs are antiviral in their native cell background. Expression of both rsIFITM isoforms significantly inhibited HCoV-229E infection in RsKT.01 cells **(Fig. 3G)**. The antiviral potency of rsIFITMs in RsKT.01 cells were comparable to that in HEK293T cells and exhibited the same pattern, with rsIFITMa showing stronger inhibition. These results indicate that while both rsIFITM isoforms are capable of restricting virus entry, they have differential antiviral specificities.

### Distinct cellular localization of *R. sinicus* IFITM isoforms

Differential antiviral activity of human IFITM1-3 can be, at least in part, explained by their distinct cellular localization (Perreira *et al*, 2013). We hypothesized that rsIFITMa and rsIFITMb likewise localize to different cellular compartments, thus influencing their ability to restrict viruses depending on their route of entry. Immunofluorescence microscopy confirmed the well-documented pattern of human IFITM1 predominating at the plasma membrane, and IFITM2/3 preferentially colocalizing with the late endosome marker CD63 **(Fig. 4A**, **Appendix figure S1)**. This localization pattern was mirrored by rsIFITM isoforms expressed in HEK293T cells – rsIFITMa in internal compartments and rsIFITMb on or near the cell surface. Surface localization was further quantified by measuring the percentage of FLAG signal at the plasma membrane, confirming that the proportion of rsIFITMb found on the cell surface was significantly higher than that of rsIFITMa **(Fig. 4B)**.

**Figure 4:**
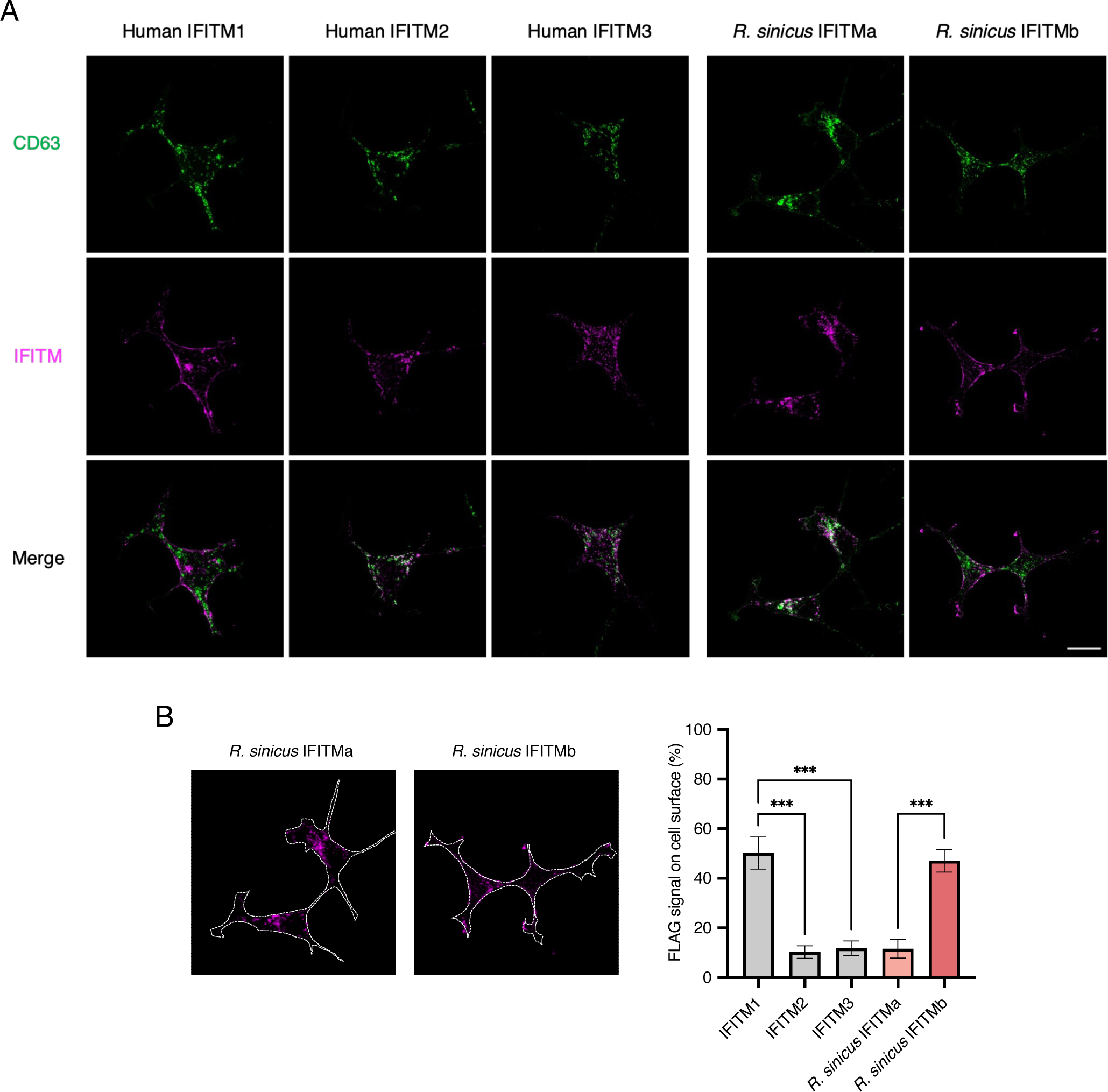
Distinct subcellular localization of *R. sinicus* IFITMs **A.** HEK293T cells were transfected with the indicated FLAG-tagged IFITMs. Cells were stained for CD63 (green; late endosome marker) and FLAG (magenta; IFITMs) at 48 hours post-transfection and imaged by confocal microscopy. Representative z-stack images are shown. Scale bar, 30 μm. **B.** FLAG signal on the surface of each cell overexpressing the indicated FLAG-tagged IFITM (dotted line) was quantified by Fiji and expressed as a percentage of the total FLAG signal from the cell. Error bars represent SEM of averages from 3-4 cells. One-way ANOVA with Bonferroni’s test; ***p<0.001.

We then examined whether the increased antiviral activity of rsIFITMa relative to rsIFITMb could be explained by its accumulation in endosomes, which may be a more favored route of entry for the studied viruses. The cathepsin L inhibitor MDL-28170 blocks the endosomal entry route of coronaviruses, forcing virus-cell membrane fusion to occur at the plasma membrane upon spike protein priming by the serine protease TMPRSS2 (Bertram *et al*, 2013). MDL-28170 therefore inhibits HCoV-229E pseudotype entry, with the inhibition being partially relieved by TMPRSS2 expression **(Fig. EV4A)**. We hypothesized that redirecting virus entry to the cell surface would enhance the antiviral potency of cell surface-localized IFITMs relative to intracellular IFITMs. Contrary to our prediction, even in the presence of MDL-28170, the predominantly endosomal rsIFITMa inhibited HCoV-229E pseudotype entry to a significantly greater extent than rsIFITMb. Similarly, the antiviral potency of human IFITM1, IFITM2 and IFITM3 relative to one another remained unchanged upon MDL-28170 treatment **(Fig EV4B)**. This suggests that the route of virus entry is insufficient to explain the difference in antiviral activities of differentially localized IFITMs.

### Evolutionary convergence of IFITM diversification strategy

Independent evolution of the IFITM family in different species has led to distinct IFITM repertoires. To understand the evolutionary relationship between IFITMs in different species, we examined the phylogeny of IFITMs from commonly studied mammals and bats of epidemiological significance. Our analysis shows that mammalian IFITMs are grouped in two ways: IFITM isoform-specific clustering and species-specific clustering **(Fig EV5A)**. Since IFITMs found in most species are not direct orthologs of human IFITMs according to their phylogenetic grouping, they were named IFITM1-, IFITM2-, or IFITM3-like based on their homology with the respective human IFITMs. Primate and rodent IFITMs cluster by isoform where IFITM1, IFITM2 and IFITM3 form separate groups, implying that these species arose after the three IFITM isoforms emerged by gene duplication. In contrast, IFITMs in all other species cluster in a species-specific manner, indicating that separation of these species occurred before the ancestral IFITM diverged independently within each species. Notably, IFITMs in bats of the suborder Yangochiroptera and Yinpterochiroptera form two distinct monophyletic groups with long branches, denoting that bat IFITMs have accumulated many mutations compared to the most recent common ancestor shared by all bat species.

Alternative splicing generates IFITM diversification as we have seen in *R. sinicus*. To understand how widespread alternative splicing is in the IFITM family, we identified *IFITM*-like genes in additional bat species from the RefSeq RNA database. Analysis was restricted to 22 bat species due to limited datasets published at the time of analysis. Species were grouped by the pattern of alternative splicing in *IFITM*-like genes they possess **(Fig EV5B)**. Alternatively spliced *IFITM*-like genes were evident in 12 bat species, but only 8 possess genes that encode non-synonymous IFITM isoforms, indicative of enhanced coding capacity **(Fig 5A)**. These bat species belong to a polyphyletic group, suggesting that they independently acquired alternatively spliced *IFITM*-like genes. For 4 of these 8 species, alternative splicing occurs by alternative first exon and results in IFITM isoforms with distinct YXXΦ endocytic motif. This subset contained *R. sinicus* alongside three other bats, including the intermediate horseshoe bat *Rhinolophus affinis*. A new reference-quality genome of *R. affinis* was recently generated through the Bat1K project, from which we identified genes showing highest homology with human immune-related *IFITM* genes (Hiller *et al*, 2023). *R. affinis* possesses two *IFITM1*-like genes, one *IFITM2*-like gene and one *IFITM3*-like gene. The *R. affinis IFITM3* gene is predicted to encode two IFITM isoforms: IFITM3a with a _20_DEML_23_-containing N-terminus, and IFITM3b with a truncated N-terminus lacking the first 21 amino acid of IFITM3a **(Appendix figure S2)**. As seen with *R. sinicus* IFITMa and IFITMb, *R. affinis* IFITM3a and IFITM3b may have distinct antiviral specificities. To confirm this, RsKT.01 cells stably expressing *R. affinis* IFITMs were challenged with HCoV-229E. All *R. affinis* IFITMs inhibited HCoV-229E infection with IFITM1-2 being the most potent inhibitor **(Fig 5B)**. *R. affinis* IFITM3a and IFITM3b indeed exhibited differential antiviral potency, where IFITM3a inhibitied HCoV-229E to a significantly greater extent. *R. affinis* therefore represents another example where IFITM functional diversity is generated by alternative splicing.

**Figure 5:**
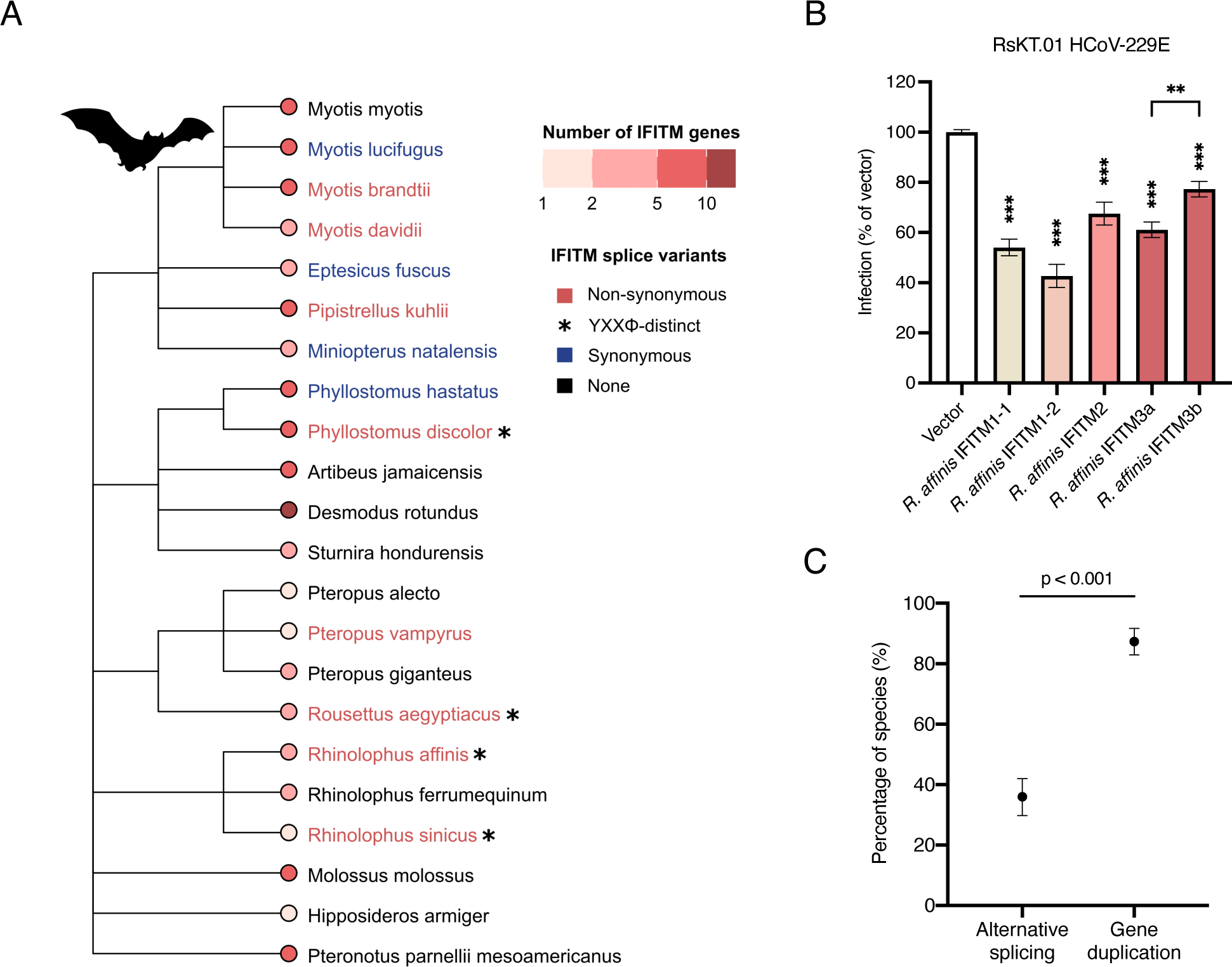
Alternative splicing generates IFITM diversity in mammals **A.** Bats were grouped according to the *IFITM*-like genes they possess. A phylogenetic tree showing the ancestral relationships between bats was labelled by their grouping: bats with *IFITM*-like gene(s) that encode two or more synonymous (blue) or non-synonymous (red) IFITMs. Bats with *IFITM*-like gene(s) encoding YXXΦ-distinct IFITM isoforms are marked with an asterisk (*). Tip nodes are coloured by the number of *IFITM-*like genes they possess. **B.** RsKT.01 cells stably expressing *R. affinis* IFITMs were infected with serially diluted HCoV-229E virus and infection was quantified by TCID50 assay. Statistical significance of difference between vector- and IFITM-expressing cells were determined by one-way ANOVA with Dunnett’s test; Statistical significance of difference between indicated groups were determined by unpaired t-test; **p<0.01, ***p<0.001. **C.** Analysis of IFITM repertoires in 206 mammalian species revealed the frequency of alternative splicing and gene duplication of *IFITM*-like genes. Error bars represent the 95% confidence interval and statistical significance of difference was determined by bootstrapping with 1000 bootstraps.

Finally, we extended the bioinformatic analysis to include all mammalian species in the RefSeq RNA database, leading to the identification of *IFITM*-like genes in an additional 184 species **(Appendix figure S3)**. Only about 5% (11/206) of these mammals have *IFITM*-like genes that encode YXXΦ-distinct IFITM isoforms, including dog, ferret, meerkat, warthog, gelada baboon, brown bear and Angolan colobus monkey. Intriguingly, 12% (25/206) of sampled mammals have only one predicted IFITM transcript. We then compared alternative splicing with gene duplication as a strategy for IFITM diversification. Gene duplication is defined to have occurred in species that possess more than one *IFITM*-like gene, and it is not mutually exclusive with alternative splicing. *IFITM* gene duplication was observed in the majority of species and is more commonly adopted by mammals than alternative splicing (87% vs 36%) **(Fig 5C)**. Notably, there is a strong association between the two strategies, meaning that species with multiple *IFITM*-like genes are more likely to also have *IFITM*-like genes that encode multiple transcripts **(Fig EV5C)**. Altogether, our evolutionary analysis uncovered alternative splicing as a previously underappreciated means of generating IFITM diversity in mammals.

## Discussion

Bats are natural reservoir hosts of pathogenic viruses with zoonotic potential, many of which are restricted by human IFITMs. Antiviral activity of human IFITM1-3 against these viruses has been widely studied, especially since the COVID-19 pandemic, but our understanding of IFITMs in species beyond human and mouse is limited. Our study shows that *R. sinicus*, the Chinese horseshoe bat, possesses an interferon-inducible *IFITM* gene that encodes two YXXΦ-distinct IFITM splice variants with conserved antiviral activity.

Using pseudotypes and replication-competent viruses, we show that both *R. sinicus* IFITM isoforms have conserved amphipathic helices and are capable of inhibiting virus infections. Notably, rsIFITMa was consistently more antiviral than the N-terminus truncated rsIFITMb. Distinct antiviral specificity is evident in the human IFITM family, with IFITM3 being the strongest inhibitor of IAV, whereas Marburg and Ebola filoviruses are more profoundly inhibited by IFITM1 (Huang *et al*., 2011). This is thought to be explained by the different localization of IFITM1 and IFITM3, and the preferred entry routes of these viruses. We therefore predicted that a similar dynamic could underlie the differential antiviral potency observed between rsIFITMa and rsIFITMb against the studied viruses. We confirmed by fluorescence microscopy that rsIFITMa and rsIFITMb indeed localize to distinct cellular compartments. rsIFITMa with a YXXΦ-containing N-terminus co-localized with the late endosomal marker CD63, consistent with the localization of microbat IFITM3 (Benfield *et al*, 2015). On the other hand, the surface localization of rsIFITMb closely resembles that observed for an N-terminally truncated murine IFITM3 which acts as a model for the human IFITM3 rs1225-C allele (Xie *et al*, 2023b). However, redirection of HCoV-229E pseudotype entry to the cell surface did not render the virus more susceptible to inhibition by rsIFITMb relative to rsIFITMa. This suggests that differences in localization relative to the site of virus entry is not sufficient to explain the distinct antiviral specificities of *R. sinicus* IFITMs. Additional factors such as expression level, post-translational modifications and interactions with virus envelopes, cellular co-factors, or other antiviral effectors may also influence antiviral specificity and potency of IFITMs. Benfield et al. previously reported that mutation in codon 70 of microbat IFITM3 altered its localization and impaired its antiviral activity (Benfield *et al*., 2020). Intriguingly, codon 70 of *rsIFITMa* and *rsIFITMb* encodes a proline and a threonine respectively, possibly contributing to their functional differences.

Our phylogenetic analysis shows that in most cases, the ancestral *IFITM* gene was duplicated after speciation, so IFITMs in different species are not orthologs of one another. We further present here, with horseshoe bats as an exemplar, that alternative splicing of *IFITM*-like genes is another means of generating IFITM diversity. Alternative splicing in the IFITM family has not been widely reported apart from the IFITM3 rs12252-C allele and the N-terminus truncated IFITM2 isoform found in immune cells. Nevertheless, our splicing analysis reveals that Chinese horseshoe bats are not unique in using alternative splicing to generate IFITM diversity. This strategy is less commonly adopted than gene duplication in mammals but is still a significant contributor towards IFITM diversity. We also found that some mammals only express one IFITM transcript, which could imply that their *IFITM* family has not evolved by either mechanism but might also result from caveats of our analysis as it is dependent on the quality of genome assemblies and RNA sequencing datasets. It is possible that limited IFITM diversity in some species is compensated by other antiviral effectors, or rich post-translational modifications that alter IFITM functions.

Alternative splicing is a source of phenotypic diversity in proteins beyond IFITMs, for example, the two splice variants of zinc finger antiviral protein (ZAPL and ZAPS) inhibit IAV infection via distinct mechanisms (Tang *et al*, 2017). An unaddressed question is whether alternative splicing always leads to functional diversity (Wright *et al*, 2022). Alternative first exon splicing regulates subcellular distribution of IFITMs and other proteins (Kim & Gladyshev, 2006). However, not all IFITM splice variants have distinct N-terminal domains as seen in Chinese horseshoe bats and intermediate horseshoe bats. Functional characterization of IFITM splice variants in other mammals is necessary to understand the contribution of alternative splicing to functional diversification of the IFITM family. It is also of interest to identify factors that influence expression levels of different gene transcripts at both the transcriptional and post-transcriptional level. For instance, herpes simplex virus-1 evades immune restriction by favoring expression of a splice variant of the antiviral myxovirus resistance protein 1 (MxA) that lacks antiviral activity (Ku *et al*, 2011). Reservoir species may possibly take advantage of IFITM alternative splicing to restrict infections in core tissues while allowing infection in others, enabling them to carry viruses asymptomatically. In the case of Chinese horseshoe bats, we speculate that endogenous expression of rsIFITMb is sufficient provide basal level protection against viruses they host, whereas the more antiviral rsIFITMa is upregulated if viral load exceeds a certain threshold to prevent disease.

Gene duplication and alternative splicing within the *IFITM* family are two strategies adopted by mammals for IFITM diversification – an example of convergent evolution. For most mammals, the use of one or more strategies to expand the IFITM toolkit highlights the importance of IFITM diversity as a component of innate immunity. Extending our splicing analysis and functional studies to other ISGs will take us beyond the genomic level to reveal distinct antiviral transcriptomes in different species – the result of evolution shaped by the virus-host arms race and the basis of species-specific zoonotic barriers.

## Materials and methods

### Identification of mammalian *IFITM* genes

*IFITM*-like genes in non-human species were identified by tBLASTn searches against the National Center for Biotechnology Information (NCBI) RefSeq RNA database using human IFITM3 as query (Boratyn *et al*, 2013). The search was restricted to mammals (Taxonomy ID: 40674) and e-value cut-off was set to 1×10^-20^ to exclude non-immune related *IFITM5*- and *IFITM10*-like genes. Non-coding RNAs were also excluded. *R. sinicus* genome with NCBI RefSeq accession GCF_001888835.1 was used. Percentage identity between *R. sinicus* IFITMs were calculated using the pairwise sequence alignment tool EMBOSS Needle (Madeira *et al*, 2022).

### RNA sequencing and analysis

RsKT.01 cells were seeded onto 6-well plates and either untreated, treated with 100ng/ml HMW poly(I:C) (Invitrogen) for 6 hours or infected with VSV at MOI=0.1 overnight. Cell lysates were collected for RNA extraction using the HP Total RNA kit (Omega Biotek). Ribosomal RNA was removed with Ribo-Zero rRNA depletion kit (Illumina) prior to cDNA synthesis using random hexamer primers. Illumina next-generation sequencing was performed on the Novaseq 6000 system (2 x 250bp) by Novogene. Raw data were trimmed and assessed for quality using FastQC v0.11.8 (Andrews, 2010). Reads were then mapped to the *R. sinicus* genome (taxid:89399) with STAR v2.7.2b (Dobin *et al*, 2013). Transcript abundances were quantified by RSEM and raw counts below 20 were removed (Li & Dewey, 2011). Differential expression analysis was performed in DESeq v1.36.0 and batch effect was corrected (Love *et al*, 2014). Cut-offs for differentially expressed transcripts were set to fold-change > 2 and p-value <0.05.

### Structural characterization of peptides

Helical wheel projection plots of IFITM amphipathic helices were generated using the HELIQUEST software (Gautier *et al*, 2008). Amphipathic helix peptides were synthesized by Vivitide and reconstituted in DMSO to produce 4mM stocks. For circular dichroism analysis, peptides were lyophilized and resuspended in 10 mM sodium borate (pH 7.4), 50mM NaCl, 25mM SDS, 3.3% ethanol and 50mM NaCl at a final concentration of 170μM. Spectra were acquired between 190 and 260nm with continuous scanning at a rate of 20nm/min on a Jasco J-1500 CD Spectropolarimeter. Spectra were recorded at 0.5nm data pitch, 1nm bandwidth and a digital integration time of 4 seconds. Secondary structure compositions of the peptides were determined using the BeStSel webserver (Micsonai *et al*, 2022).

### NBD-cholesterol binding assay

Binding of amphipathic helix peptides to NBD-cholesterol was assessed as described previously (Rahman et al., 2022). In brief, 500nm NBD-cholesterol (Thermo Fisher, N1148) was incubated with serial dilutions of peptides (0-100μM) in black-wall clear-bottomed 96-well plates. Plates were incubated for 1 hour at room temperature or 4°C before measuring fluorescence intensity and polarization respectively. Measurements were taken by a Tecan Infinite M1000 at 470nm excitation and 540nm emission. A rotavirus NSP4-derived peptide was used as negative control.

### Cell culture

HEK293T (ATCC and ECACC), HEK293T-ACE2-TMPRSS2 (NIBSC), Huh7.5 cells (gift from Jürgen Haas) and RsKT.01 cells were cultured in DMEM with GlutaMAX (Thermo Fisher), supplemented with 10% fetal bovine serum and 1% penicillin-streptomycin (Gibco) at 37°C and 5% CO_2_. Media was supplemented with non-essential amino acids for Huh7.5 cells and normocin (Invitrogen) for RsKT.01 cells.

### Transfection and western blotting

rsIFITMa and rsIFITMb constructs with a FLAG tag at the N-terminus were synthesized by IDT and cloned into the pQCXIP vector for transfection. HEK293T, Huh7.5 cells were transfected with Lipofectamine 2000 (Invitrogen) and FuGENE HD (Promega) reagent respectively. At 48 hours post-transfection, cells were lysed in 1% Triton X-100 buffer with phosphatase and protease inhibitors (Roche). Protein was separated by 4-12% Bis-Tris protein gel (Bio-Rad) and transferred to 0.2 μm PVDF membrane (Cytiva). The following antibodies were used: mouse anti-FLAG-M2 (Sigma), mouse anti-beta-actin (Santa Cruz Biotechnology), IRDye 800CW goat anti-mouse (LICOR), IRDye 800CW goat anti-rabbit (LICOR). Protein was detected on a Li-Cor Odyssey imaging system.

### Stable cell line production

*R. sinicus* and *R. affinis* IFITM constructs were synthesized by Tsingke Biotech and cloned into the pLVX-IRES-mCherry vector for lentivirus production by co-transfecting HEK293T cells with the psPAX2 plasmid. Lentiviral supernatant was passed through a 0.45μm filter and added to RsKT.01 cells with 4μg/ml polybrene (Biosharp) for 4-6 hours. At 72 hours post-transduction, cells were sorted for mCherry fluorescence and pooled to culture as stable cell lines.

### Replication-competent virus infection

Transfected HEK293T cells were seeded onto 24-well plates one day prior to infection with Influenza virus A/Puerto Rico/8/1934 (H1N1, PR8) at MOI=0.05 for 18 hours at 37°C. Huh7.5 cells were infected with HCoV-229E-GFP at MOI=0.05 for 18 hours at 34°C. To measure infection, cells were fixed and permeabilized with the Cytofix/Cytoperm kit (BD Biosciences), stained using 1:500 rabbit anti-FLAG (Sigma) and mouse anti-IAV NP (Abcam), followed by 1:300 Alexa Fluor 488-conjugated goat anti-mouse (Invitrogen) and Alexa Fluor 647-conjugated goat anti-rabbit (Invitrogen), and analyzed on a LSRFortessa flow cytometer (BD Biosciences). RsKT.01 cells stably expressing IFITMs were infected with serially diluted HCoV-229E in low-serum media. Infection was measured by observing cytopathic effects at 3-5 days post-infection to calculate TCID_50_ using the Spearman and Karber algorithm.

### Lentiviral-based pseudotyped viruses production and transduction

SARS-CoV-1 and MERS-CoV pseudotypes were produced by co-transfecting HEK293T cells with pNL4-3.LucR^-^E and pcDNA3.1-SARS-CoV-1-spike or pcDNA3.1-MERS-CoV-spike. HCoV-229E pseudotype was produced by co-transfecting HEK293T cells with p8.91 Gag-Pol, pCSFLW and pcDNA3.1-229E-spike plasmids (Cantoni *et al*, 2022). Supernatant was harvested at 72 hours post-transfection, passed through a 0.45μm filter and stored at −80°C. Target cells that transiently express the respective virus entry receptor were incubated with pseudotypes for 48 hours, pseudotype entry was then quantified using Bright-Glo luciferase assay (Promega). For experiments involving drug pre-treatment, target cells were incubated with DMSO or MDL-28170 (20 μM) for 1 hour prior to transduction.

### Confocal immunofluorescence microscopy

Cells transiently expressing IFITMs were seeded onto 8-well chamber slides at 50,000 cells per chamber. After 24 hours, cells were fixed and permeabilized with the Cytofix/Cytoperm kit (BD Biosciences), stained using 1:400 mouse anti-CD63 (Santa Cruz Biotechnology) and rabbit anti-FLAG (Sigma), followed by 1:300 Fluor 488-conjugated goat anti-mouse (Invitrogen) and Alexa Fluor 647-conjugated goat anti-rabbit (Invitrogen). Slides were mounted with ProLong glass antifade mountant (Invitrogen) and imaged on the Leica TCS SP8 confocal microscope. Z-stack processing and further analyses were performed in Fiji (Schindelin *et al*, 2012).

### Construction of IFITM phylogenetic tree

Phylogenetic analysis of mammalian IFITMs was performed on MEGA X (Kumar *et al*, 2018). To exclude non-functional IFITMs, the following criteria was used for IFITM inclusion: protein length of 102-157 amino acids; contains 2 exons; presence of CD225 and a transmembrane domain (Hickford *et al*., 2012; Ikeda, 2003; Marchler-Bauer & Bryant, 2004; Marchler-Bauer *et al*, 2011). The best-fit nucleotide substitution model was determined by maximum likelihood analysis and a tree was constructed from IFITM protein-coding nucleotide sequences with 1000 bootstraps. The tree was annotated in FigTree (Rambaut, 2018).

### *IFITM* alternative splicing analysis

Alternative splicing patterns of mammalian *IFITM-*like genes were characterized by analyzing mRNA transcripts produced from identified mammalian *IFITM*-like genes using NCBI Datasets (Sayers *et al*, 2021). Transcripts encoded by the same gene were compared by nucleotide alignment and genes were grouped based on their splicing pattern. Phylogenetic tree inferring the evolutionary relationships between analyzed species was generated on NCBI taxonomy browser (Schoch *et al*, 2020).

## Data availability

No primary datasets have been generated and deposited.

## Acknowledgements

We would like to thank Dr. Marjolein Kikkert (Leiden University Medical Center) for useful discussions and Dr. Kin Kui Lai (National Institutes of Health) for help with production of MERS-CoV and SARS-CoV-1 pseudotypes. We also thank Prof. Jürgen Haas (University of Edinburgh) and Prof. Jincun Zhao (Guangzhou Institue of Respiratory Health) for cells and HCoV-229E viruses. NM is funded by a joint Ph.D. studentship from the University of Edinburgh and Leiden University Medical Center. JT is funded by a Medical Research Council doctoral training program (MR/N013166/1). ATI is funded by a Key grant from the National Science Foundation of Zhejiang Province (Z23C010003) and a National Science Foundation Research Fund for International Excellent Young Scientists (RFIS-II, 82350610279).

## Author contributions

**Nelly Mak:** Conceptualization; investigation; methodology; data curation; formal analysis; software; validation; visualization; writing – original draft. **Dan Zhang:** Investigation; formal analysis. **Xiaomeng Li:** Software; formal analysis. **Kazi Rahman:** Investigation; methodology. **Siddhartha A.K. Datta:** Investigation; methodology. **Jordan Taylor:** Software, writing – review & editing. **Jingyan Liu:** Investigation. **Nigel Temperton:** Methodology; resources; writing – review & editing. **Zhengli Shi:** Resources. **Aaron T. Irving:** Conceptualization; funding acquisition; investigation; resources, writing – review & editing. **Alex A. Compton:** Conceptualization; funding acquisition; resources; supervision; writing – review & editing. **Richard D. Sloan:** Conceptualization; funding acquisition; project administration; supervision; writing – review & editing

## Expanded view figure legends

**Expanded view figure 1:**
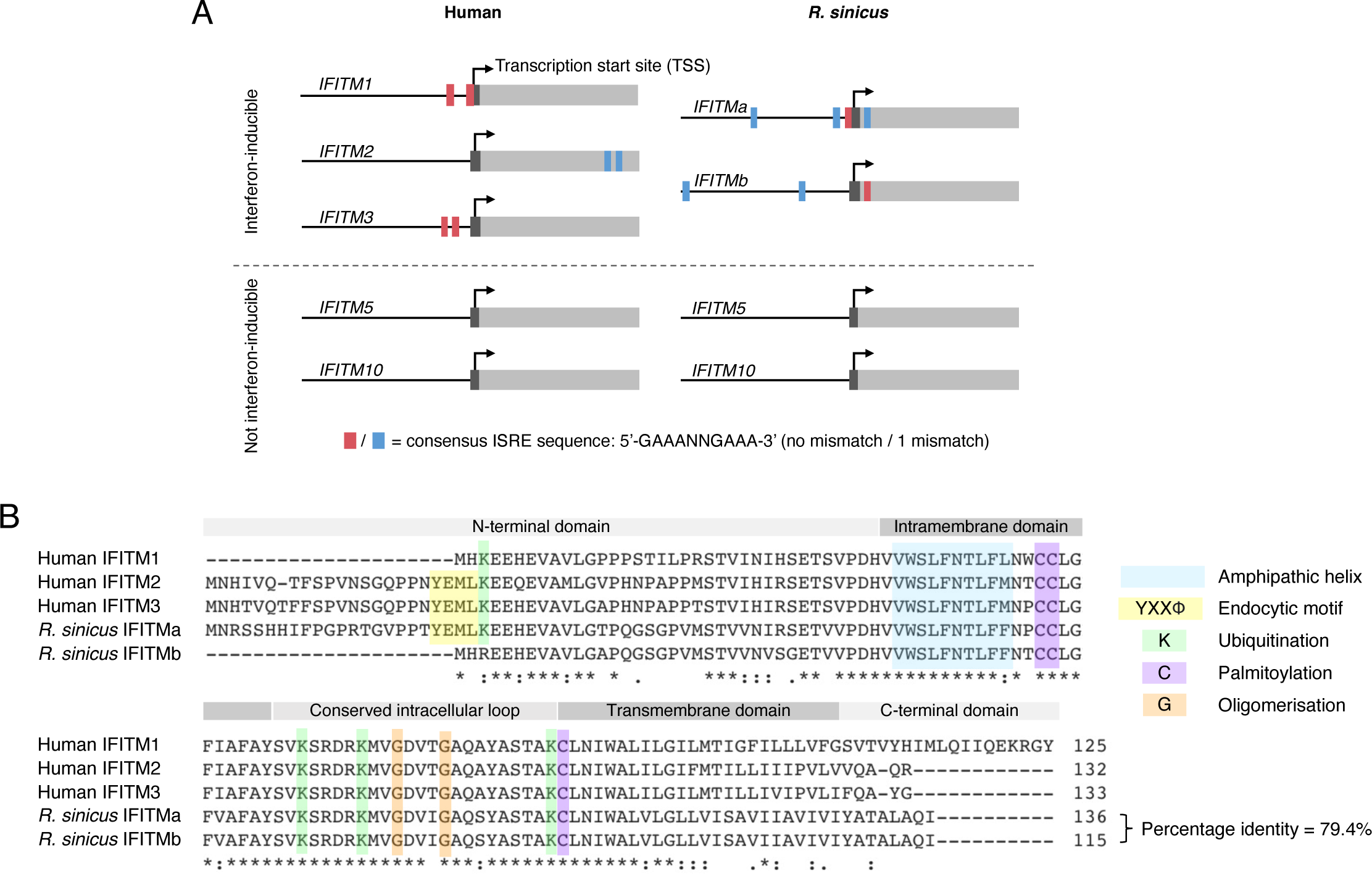
Alignment and identification of interferon-stimulated response elements (ISRE) in human and *R. sinicus* IFITMs **A.** Identification of ISRE motifs around the transcription start site (± 350 base pairs) of human and *R. sinicus IFITM* genes. ISREs were defined to have the consensus sequence of GAAANNGAAA or TTTCNNTTTC, with no mismatch (red) or 1 mismatch (blue) (Lee *et al*, 2018). **B.** Amino acid sequence alignment of immune-related human and *R. sinicus* IFITMs. Protein domains, functional motifs and amino acids that undergo post-translational modifications are highlighted. Asterisks (*) indicate positions with a conserved residue; colons (:) and periods (.) indicate conservation between groups of strongly and weakly similar properties respectively. Percentage identity was calculated using the pairwise sequence alignment tool EMBOSS Needle (Madeira *et al*., 2022).

**Expanded view figure 2:**
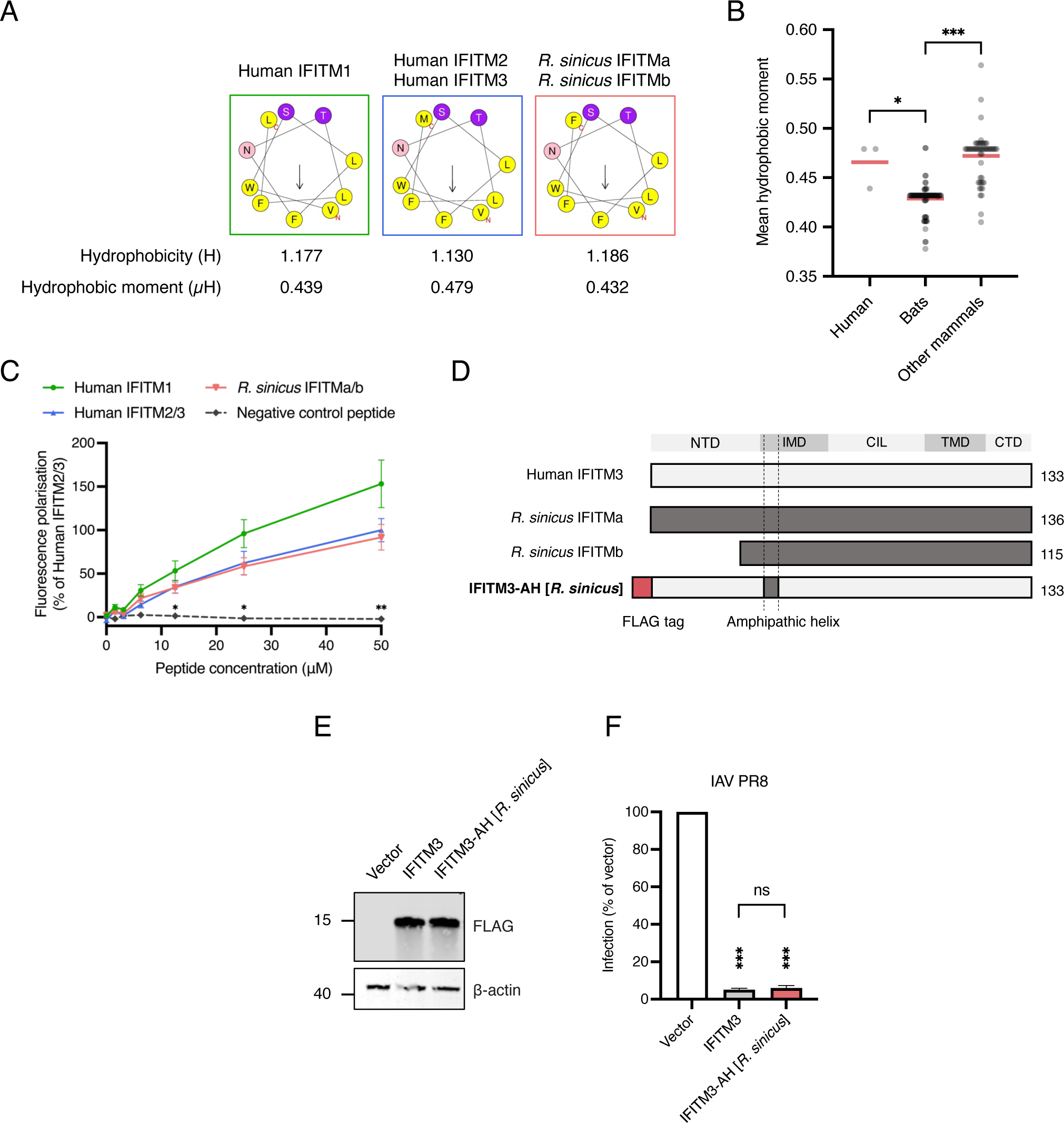
Characterization of *R. sinicus* IFITM amphipathic helix **A.** Helical wheel project plots of human and *R. sinicus* IFITM amphipathic helices containing hydrophobic (yellow) and hydrophilic (purple or pink) residues. Arrows indicate magnitude and direction of mean hydrophobic moments. **B.** Mean hydrophobic moment of IFITM amphipathic helices from 34 mammalian species, including 19 bat species. Medians of each group are shown. Kruskal-Wallis test; *p<0.05, ***p<0.001. **C.** NBD-cholesterol fluorescence polarization was measured following incubation of IFITM amphipathic helix peptides (0-50μM) with NBD-cholesterol (500nM). Data points are normalised to 50uM human IFITM2/3. (Error bars represent SEM of averages from 3 independent experiments. Statistical significance of differences between human IFITM2/3 and another peptide was determined by one-way ANOVA with Dunnett’s test; *p<0.05, **p<0.01, ***p<0.001. **D-F.** HEK293T cells were transfected with FLAG-tagged IFITM3 or chimeric IFITM3-AH [*R. sinicus*] as illustrated in (D), and protein expression was detected by western blotting at 24 hours post-transfection (E). Transfected cells were infected with IAV and analyzed by flow cytometry at 18 hours post-infection. Error bars represent SEM of averages from 3 independent experiments, each performed in duplicate. Statistical significance of difference between vector- and IFITM-transfected cells was determined by one-way ANOVA with Dunnett’s test; Statistical significance of difference between IFITM3- and IFITM3-AH [*R. sinicus*]-transfected cells was determined by unpaired t-test; ***p<0.001; ns, non-significant.

**Expanded view figure 3:**
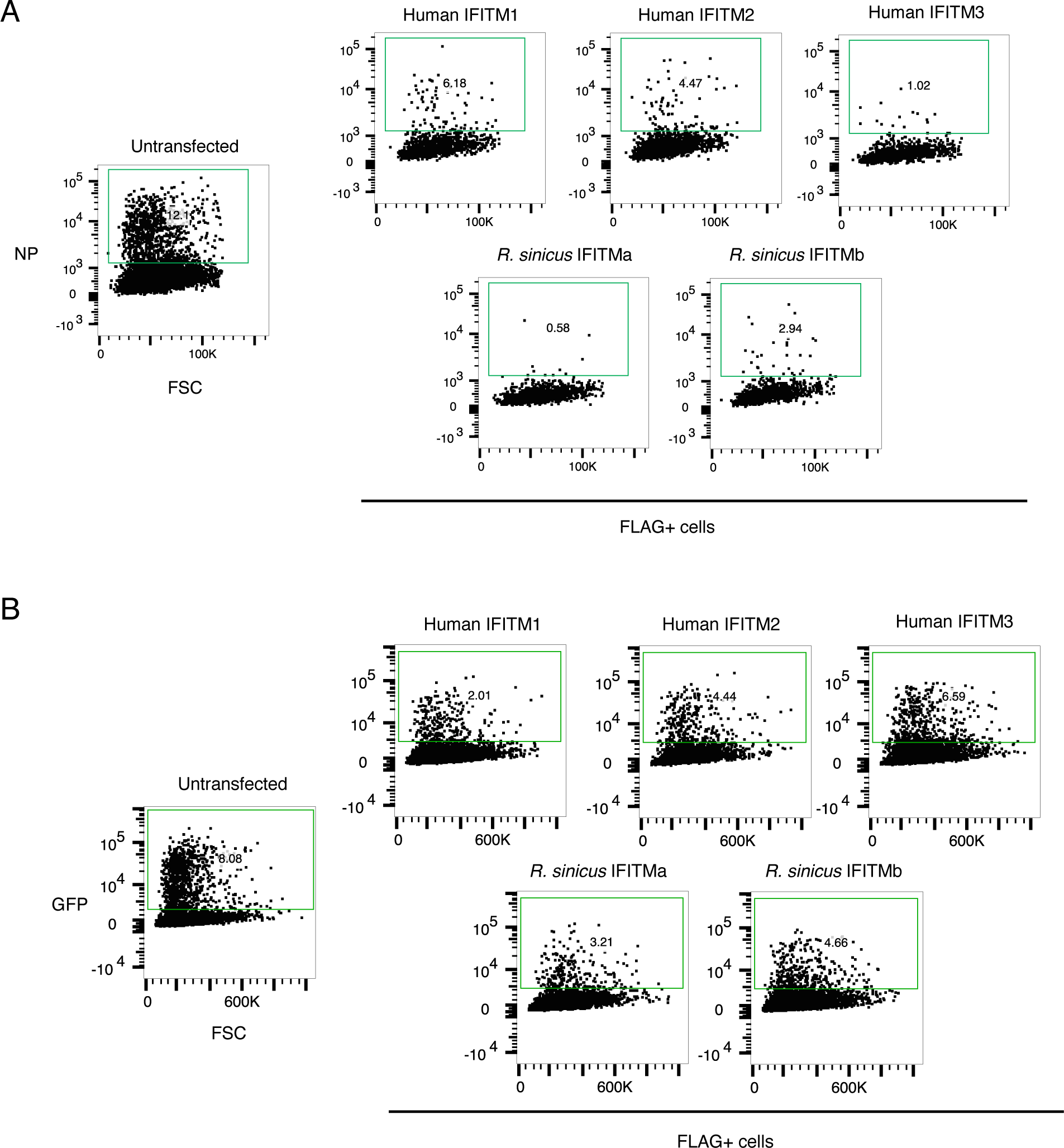
Flow cytometry analyses of virus restriction by IFITMs HEK293T **(A)** or Huh7.5 **(B)** cells were transfected with FLAG-tagged IFITM constructs and infected with IAV or HCoV-229E-GFP for 18 hours. Flow cytometry dot plots show gating for IAV NP staining or GFP signal. Transfected cells were first gated for FLAG staining and numbers indicate percentage of NP+ or GFP+ cells within the gated population. Plots are representative of three independent experiments, each performed in duplicate.

**Expanded view figure 4:**
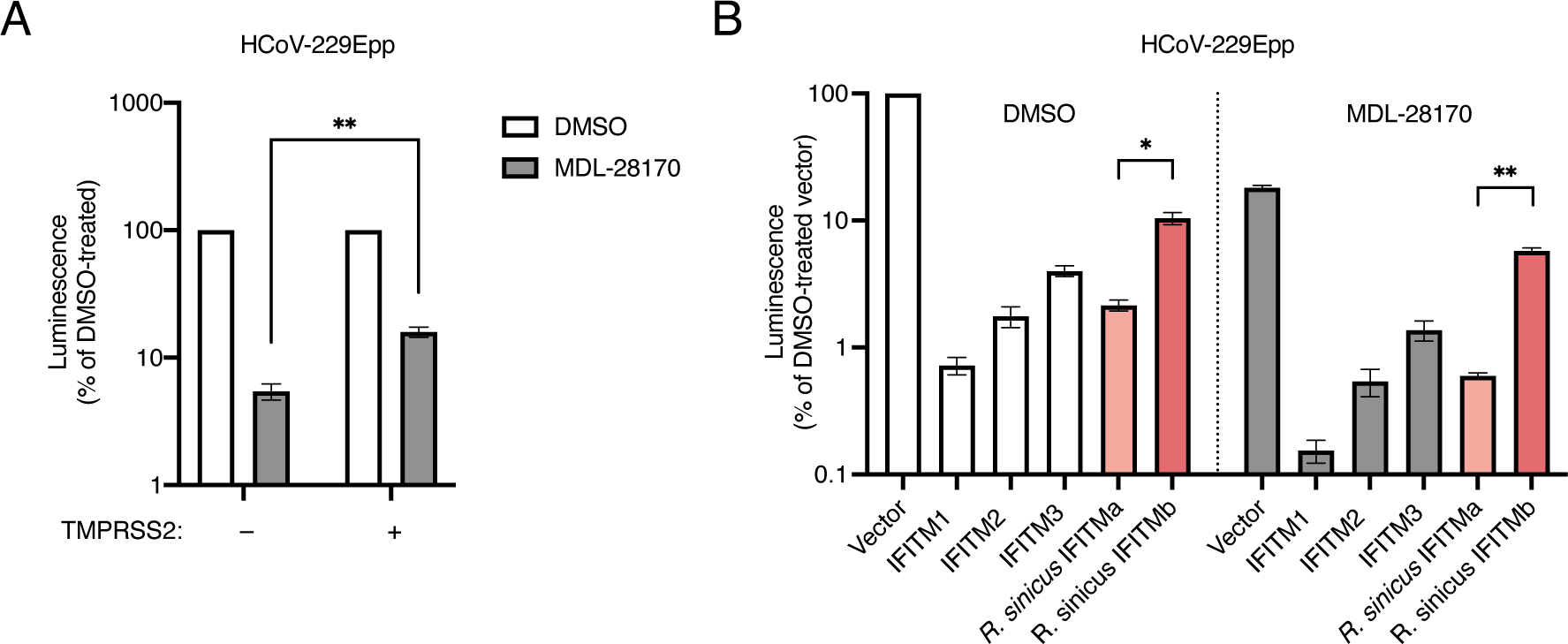
The effect of virus entry route on IFITM sensitivity **A.** HEK293T and HEK293T-ACE2-TMPRSS2 cells were transfected with APN and transduced with HCoV-229E pseudotypes in the presence of DMSO or MDL-28170. Cells were lysed and analyzed by luciferase assay after 48 hours. Data points were normalized to the respective DMSO-treated cells (white bars). Error bars represent SEM of averages of three independent experiments, each performed in triplicate. **B.** HEK293T-ACE2-TMPRSS2 cells were co-transfected with APN and the indicated FLAG-tagged IFITM constructs, then transduced with HCoV-229E pseudotypes in the presence of DMSO or MDL-28170. Data points were normalized to vector/DMSO. Error bars represent SEM of averages from 3 independent experiments, each performed in triplicate. Statistical significance of differences was determined by unpaired t-test; *p<0.05, **p<0.01.

**Expanded view figure 5:**
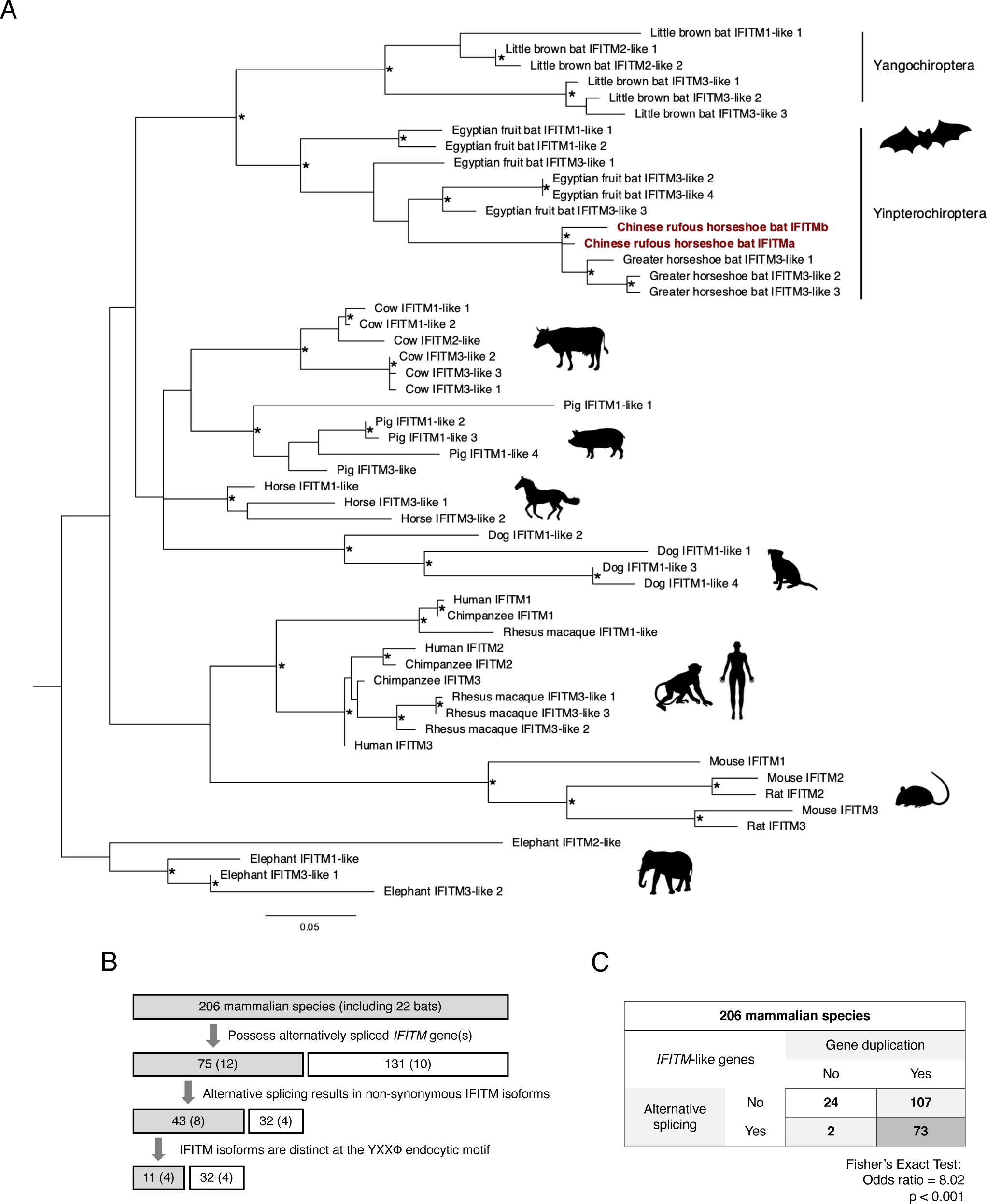
Analysis of mammalian *IFITM*-like genes **A.** Phylogenetic tree constructed by maximum likelihood analysis of 54 IFITM protein-coding nucleotide sequences from different mammalian species. The tree was rooted on the elephant outgroup and nodes with bootstrap values >70% are marked with asterisks (*). Scale bar corresponds to 0.05 substitutions per site. **B.** Flow chart illustrating the classification of *IFITM* genes by their pattern of alternative splicing in 206 mammalian species, including 22 bats. **C.** Association between alternative splicing and gene duplication in the *IFITM* family was tested by the Fisher’s exact test.

## Appendix

**Appendix figure S1:**
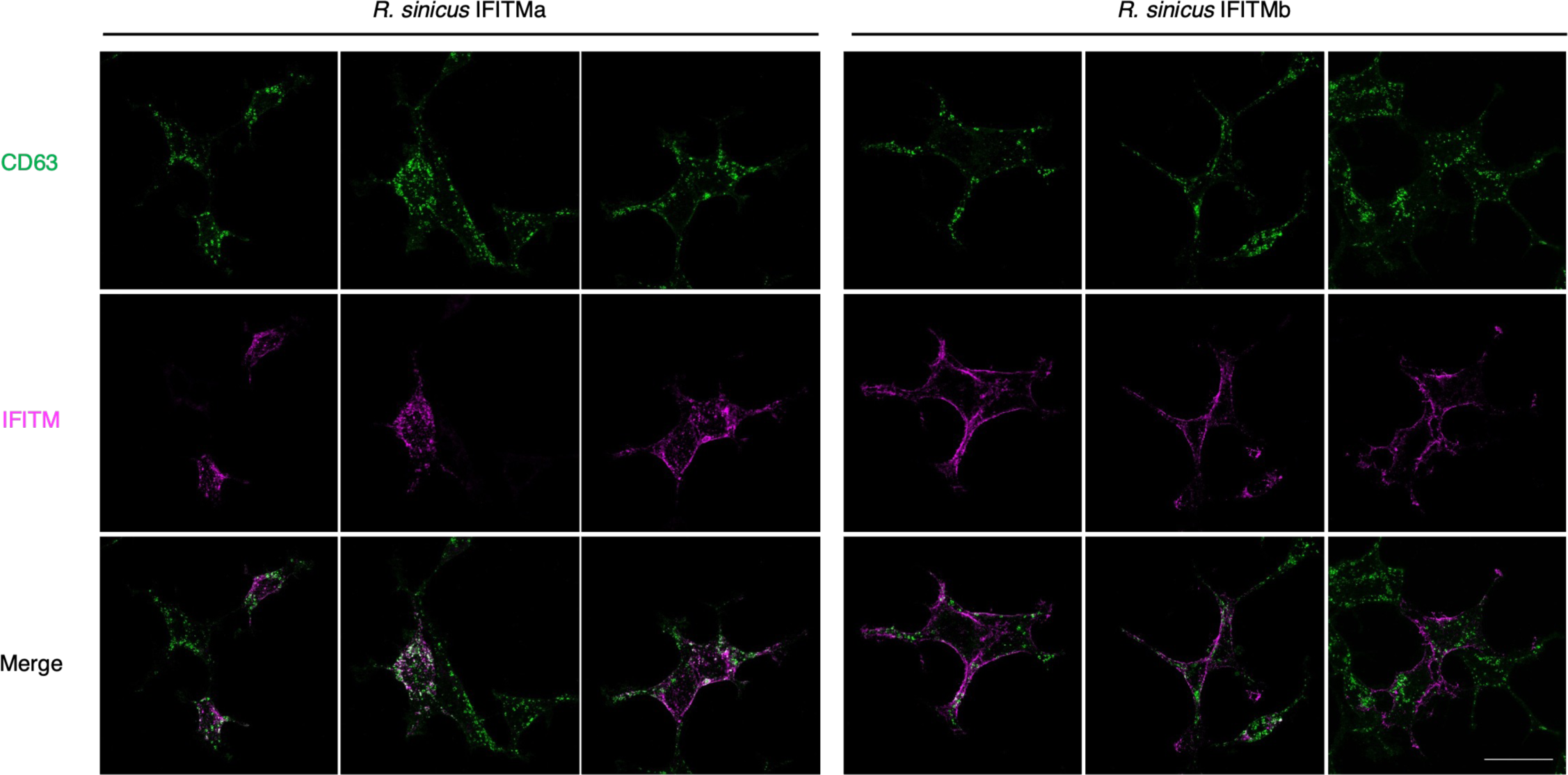
Expanded panel of immunofluorescence images of *R. sinicus* IFITMs HEK293T cells were transfected with FLAG-tagged *R. sinicus* IFITMa or IFITMb. Cells were stained for CD63 (green; late endosome marker) and FLAG (magenta; IFITMs) at 48 hours post-transfection and imaged by confocal microscopy. Representative z-stack images are shown. Scale bar, 30μm.

**Appendix figure S2:**
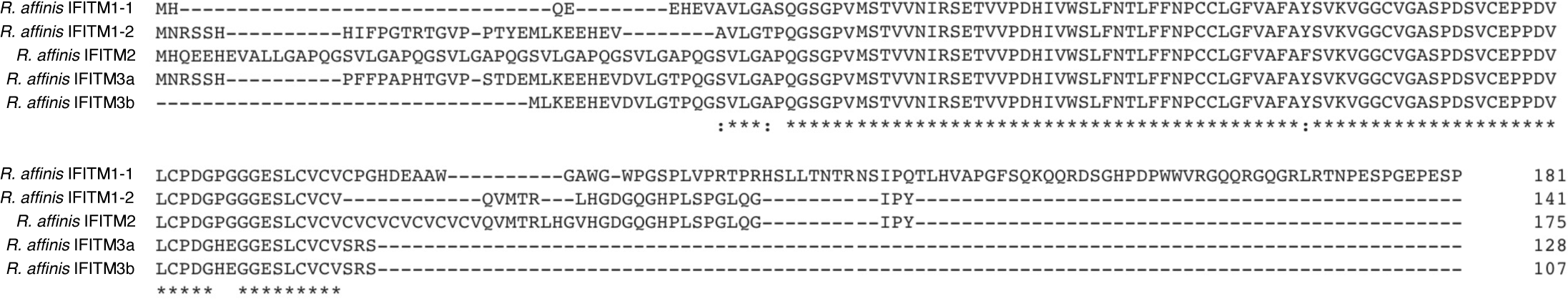
Sequence alignment of *R. affinis* IFITMs Protein sequence alignment of *R. affinis* IFITMs that show highest homology with human immune-related IFITMs. Asterisks (*) indicate positions with a conserved residue; colons (:) and periods (.) indicate conservation between groups of strongly and weakly similar properties respectively.

**Appendix figure S3:**
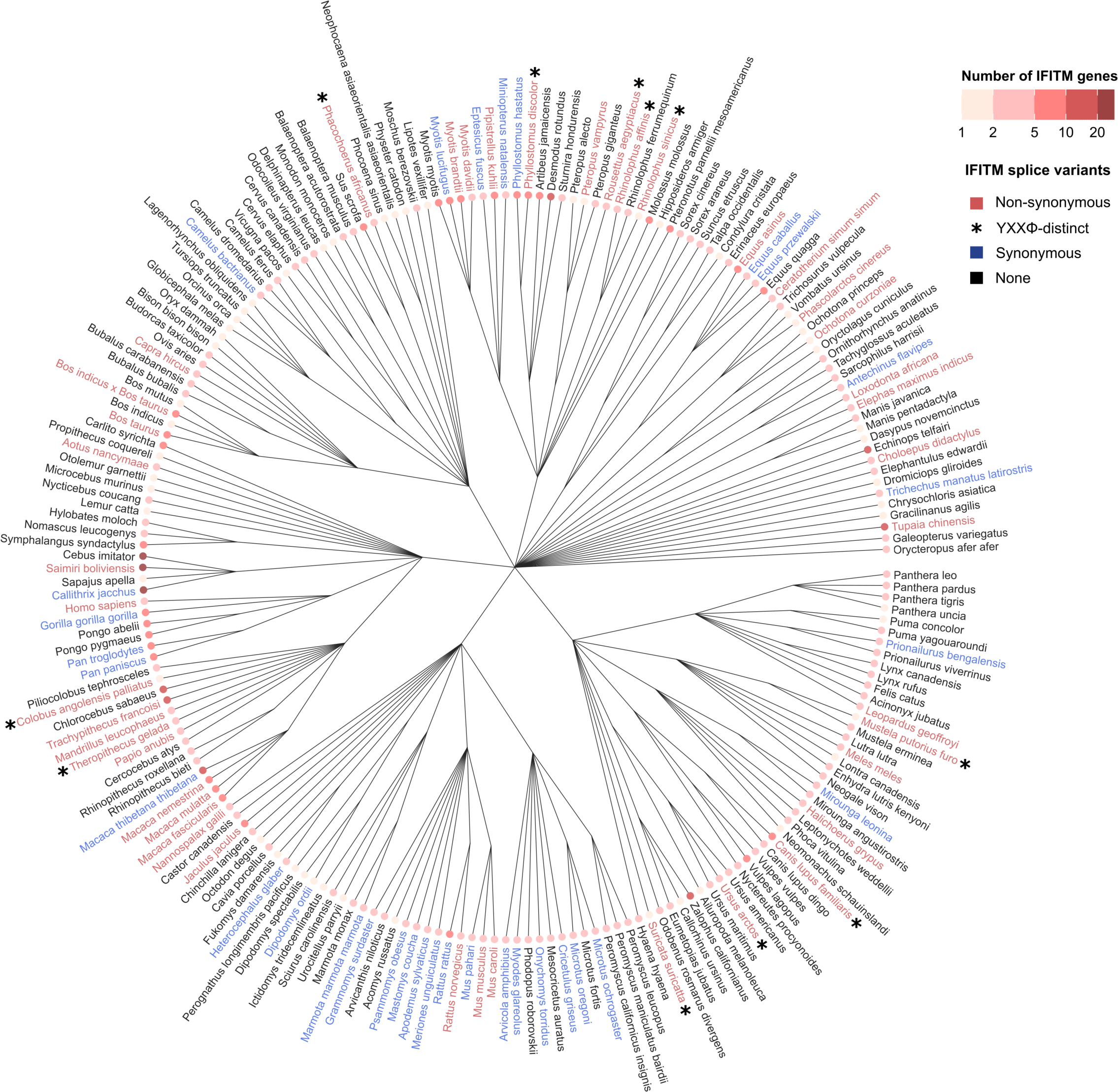
Extended splicing analysis of mammalian *IFITM*-like genes Analysis of *IFITM*-like genes in Figure 5A is extended to include 205 mammalian species. Mammals were grouped according to the *IFITM*-like genes they possess. Phylogenetic tree showing the ancestral relationships between these species was labelled by their grouping: species with *IFITM*-like gene(s) that encode two or more synonymous (blue) or non-synonymous (red) IFITMs. Species with *IFITM*-like gene(s) encoding YXXΦ-distinct IFITM isoforms are marked with an asterisk (*). Tip nodes are coloured by the number of *IFITM-*like genes they possess.

